# The use of autologous skeletal muscle derived cells as a sling in the treatment of induced stress urinary incontinence, an experimental study

**DOI:** 10.1101/2021.06.15.448536

**Authors:** Bassem S. Wadie, Haytham G. Aamer, Sherry M. Khater, Mahmoud M. Gabr

## Abstract

**Introduction & hypothesis:** This is an experimental pre-clinical study testing for the applicability of autologous skeletal muscle derived cells as a seeded sling for the treatment of SUI in the canine model

**Methods:** 10 Mongrel dogs: In 4, skeletal muscle biopsy was harvested from Biceps Femoris. 1 month later, incontinence was induced in 8 dogs through surgical disruption of the pubourethral ligaments. Muscle biopsy was incubated in DMEM medium and after expansion for 8 weeks, MDCs were collected. PGA scaffold was immersed in culture medium, coated with matrigel and cells were seeded. The sling was placed suburethrally in 8 dogs; 2 of which were cell-seeded and 4 had the scaffold only.

Urethral pressure (UP) measurement was done at baseline and 2 weeks alter insertion of the sling.

The urethra with its surrounding was harvested 4 weeks after sling insertion for histopathology. 2 dogs were considered as control, in which no urethrolysis or insertion of slings were carried out

**Results:** UP show increase of maximum urethral pressure during static measurement in all dogs with a scaffold inserted. The increase ranged from 5-40 cmH20 (Median 23 cmH20)

Histopathology shows significant periurethral proliferation of skeletal muscles in 4 dogs with cell-seeded scaffold, as demonstrated by Desmin. This was maximum in dogs # 1& 2. This was not the case in the 4 dogs that had PGA sling only.

**Conclusion:** The use of skeletal muscle –seeded PGA scaffold is a practical technique with preserved integrity of histological differentiation in canine model at short term.

**Brief Summary:** Autologous Skeletal muscle-derived cells could be propagated in vitro and seeded to PGS scaffolds and used as slings

## Introduction

Treatment of stress incontinence is challenging^1^. Different techniques evolved throughout clinical trials^2^. So far, no procedure could be considered “gold”. However, a midurethral sling (MUS) is the most widely used procedure and is considered as the standard of care in women with stress incontinence^3^. Synthetic MUS are not without risks and sometimes could be life-threatening^4^. Many trials, both experimental and human, attempted to provide an autologous, efficacious and durable tissue-engineered sling.

This study evaluates a biodegradable *seeded Polyglcolic acid (PGA) scaffold* enriched with autologous muscles, so as to produce a sling without the need for a harvesting procedure such as that implemented in rectus fascia sling ^5^, being the prototype of autologous slings^6^.

## Materials and methods

The study comprised 10 Mongrel female dogs. Approval from the local Ethics Committee was obtained. Open episiotomy was carried out 2 weeks prior to the study to facilitate the vaginal approach.

Group 1 comprises 4 dogs in which a cell-seeded sling was applied while in group 2, only the PGA sling was applied.

### Isolation and expansion of muscle-derived cells (MDCs)

In 4 dogs (group 1), muscle biopsy was harvested from biceps under general anesthesia. Obtained biopsy averaged 0.2 gm and was incubated in *0.2% collagenase type 1A in* Dulbecco’s Modified Eagle Medium *(DMEM) medium for 20 minutes at 37 C*. (Sigma-Aldrich®, St. louis, USA) digestion. Cells were cultured on laminin-coated tissue culture flasks in SKGM-2 medium (Lonza®, Valkersville, MD). Medium was replaced every 3 days. At confluence, cells were passaged and split 1:2 in tissue culture flasks. After 8 weeks and 4 passages, MDCs were collected for transplantation.

- **Characterization of MDCs:** Morphology of cells was evaluated using phase-contrast microscopy. The expression of Desmin was assessed by immunohistochemistry (IHC), using monoclonal anti-Desmin antibodies (DakoCytomation®, Glostrub, Denmark).
- **Cell seeding on scaffolds:** Neoveil absorbable PGA sheet (Gunze Limited®, Kyoto, Japan) was used as (2×3 cm) segments. Seeding started by immersion in culture medium supplemented with 10 % fetal bovine serum and 1 % penicillin/streptomycin for 24 h. at 37 ° C before seeding. The scaffold was coated with matrigel and MDCs were seeded at a concentration of 1 million cells/ cm^2^. After 24 h. the seeding side was flipped and the other side was treated the same way. Both surfaces were fully immersed in medium during seeding process. Seeded scaffold was cultured for further 3 days in the same culture medium
- **Induction of incontinence:** One month after biopsy, incontinence was induced in 8 dogs through midline suprapubic incision and identification of the urethra and bladder. Sharp dissection of the urethra all around until disruption of the periurethral sphincter is achieved. This method is similar to that previously described by Rodriguez et al^7^, with the addition of pubourethral ligament disruption.
- Two weeks after induction of incontinence, sling was applied through vaginal incision in suburethral position in all 8 dogs: Cell-seeded scaffold in the first 4 and scaffold – only in the other 4. The sling is fixed to the periurethral tissue. Urethral pressure (UP) measurement was done before (baseline) and 2 weeks after insertion of the sling.
- The urethra with its surrounding was harvested and dogs were sacrificed 4 weeks after sling insertion. Figure 1 shows the scaffold and steps of surgical procedure
- The urethra was fixed in 10 % buffered formalin and processed in paraffin blocks, sectioned at 5 μm and stained with hematoxylin and eosin (H&E), methenamine silver (for reticuline fibers detection) and Masson trichrome stain (for fibrosis detection and muscle bundles delineation).
- 2 dogs were considered as control, in whom no urethrolysis or insertion of slings were carried out. UP was carried out in these dogs before being sacrificed and their urethra was taken as control.
- For IHC, desmin antibodies were used (Abcam, Cambridge, UK) at dilution of 1:100. Nerve fibers were identified through staining with polyclonal anti-S100 (DAKO, Carpinteria, CA) at a dilution of 1:200. Immunolabeling was performed using an avidin–biotin detection kit (Vectastain Elite ABC, Vector, Bulingame, CA). All sections were counterstained with Gill’s hematoxylin.

**Figure 1:**
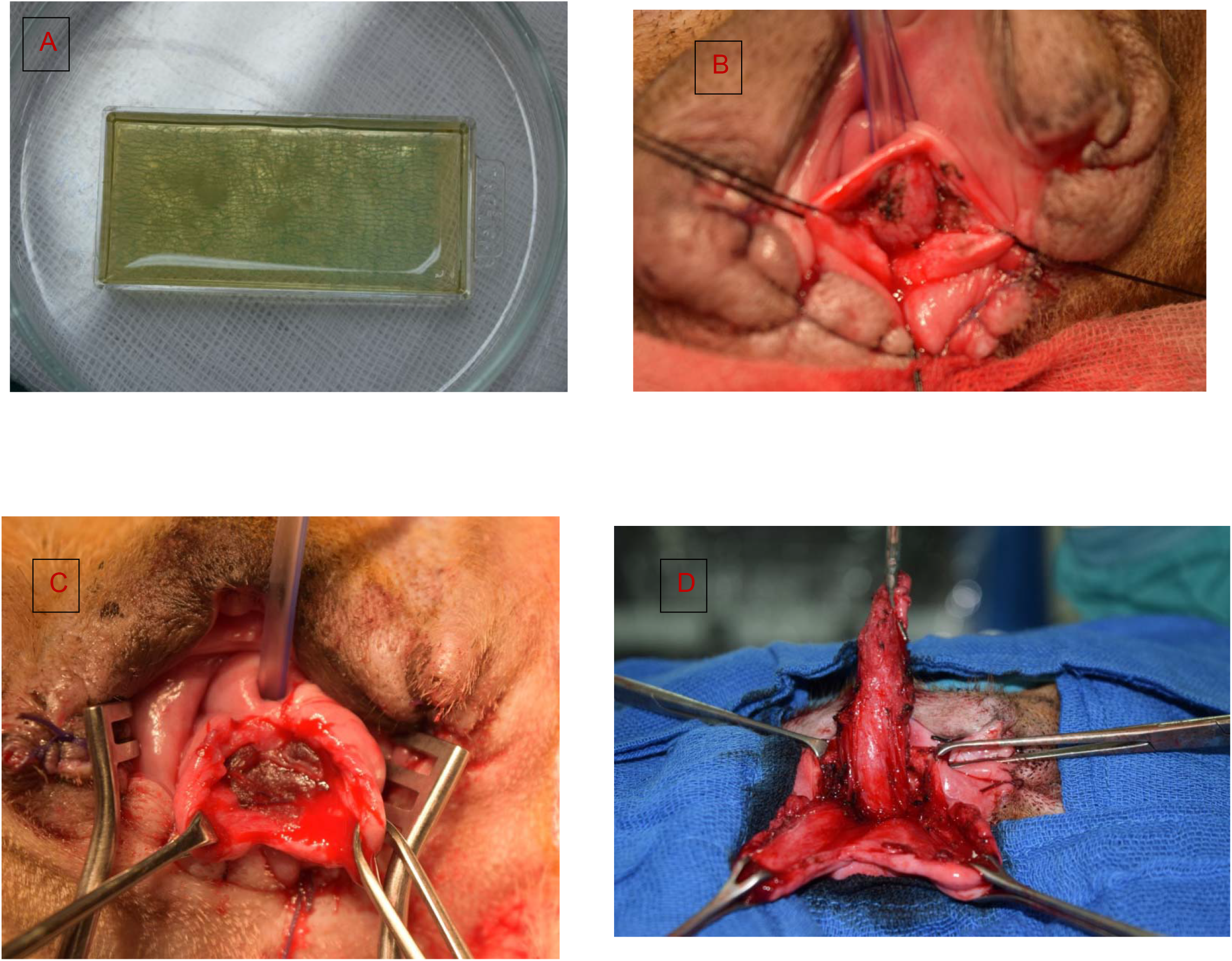
A) Double -faced seeded scaffold in sterile medium. B)Midline vaginal incision with distal 2/3 of the dog urethra completely exposed, c) PGA sling is attached to the periurethral tissue using 5/0 Vicryl, D) Whole urethra is dissected from the surrounding, and harvested

A flowchart for the research procedures was supplemented (Supplement)

## Results

- UP shows increase of maximum urethral pressure in all dogs with a scaffold inserted. Table 1 demonstrates change in UP over time. The increase in maximum urethral closure pressure (MUCP) ranged from 12-60 cmH20 (Median 40.25 cmH20) in the first 4 dogs, where the scaffold was seeded with skeletal muscle cells. In the scaffold -only group, the increase of MUCP ranged from −3 to 18 (median 5.5 cmH20)
- Histopathology: Luminal diameter, lining epithelium, lamina propria thickness with its elastic and collagenous fibers content (normally about 70% of whole sphincter thickness), fibrous tissue deposition and thick walled veins condition (stratum cavernosum), smooth and striated muscle(both constituting about 30% of total urethral thickness)^8^ were assessed.

**Table 1:**
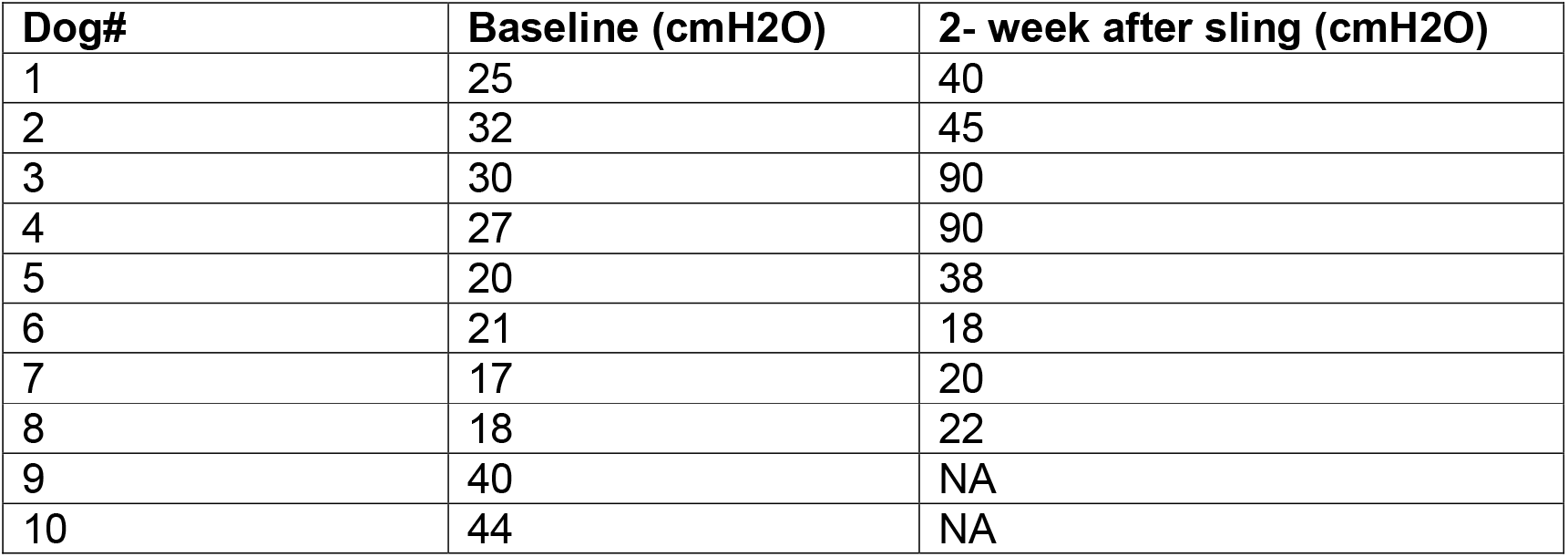
Maximum urethral closure pressure (MUCP) in all dogs.

| **Dog#** | **Baseline (cmH2O)** | **2- week after sling (cmH2O)** |
| --- | --- | --- |
| 1 | 25 | 40 |
| 2 | 32 | 45 |
| 3 | 30 | 90 |
| 4 | 27 | 90 |
| 5 | 20 | 38 |
| 6 | 21 | 18 |
| 7 | 17 | 20 |
| 8 | 18 | 22 |
| 9 | 40 | NA |
| 10 | 44 | NA |

### In group 1

urethral lumen was slightly dilated in comparison to the control group; mucosal lining is of near normal thickness. Lamina propria is slightly thicker and no evidence of fibrous tissue deposition as evidenced with Masson trichrome. Reticular fibres are the same as in control group. Thick walled veins are mildly increased in number and caliber compared to control group. The overall muscle layer appears as lavender red bundles in Masson trichrome are normal in thickness with some degree of crowding and disorganization. Areas of newly formed muscle bundles with central nuclei were confirmed.

### In group 2 (scaffold only)

The damaged sphincter shows mildly dilated luminal area, with atrophic urothelial lining. The lamina propria thickness is increased with moderate fibrous tissue deposition in addition to lower reticular fiber content and mild inflammatory lymphocytic infiltrate. The vascular component is unremarkable. There is substantial loss of muscle layer with an overall decreased wall thickness.

IHC for anti-desmin shows significantly higher sphincter muscle mass in seeded scaffold specimens as compared to the other group. Newly formed muscle bundles are wider and more randomly aligned, compared to normal sphincter. Staining for S100 shows the presence of nerve fibers between the regenerated sphincter muscles. Figure 2 and 3 shows histopathological changes in the two groups.

**Figure 2:**
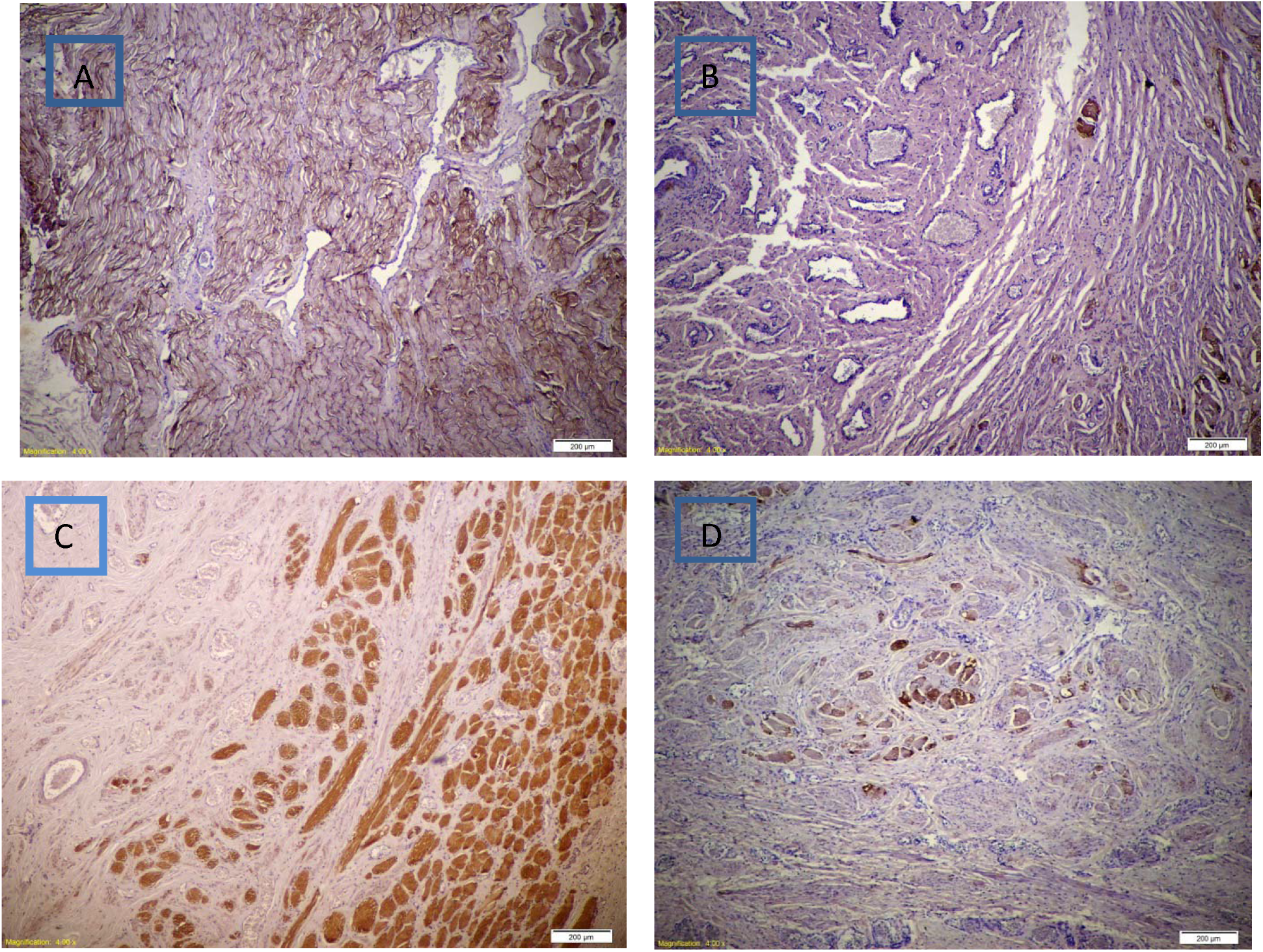
A} Diffuse increase in thickness of skeletal muscle bundles with mild disorganization in PGA seeded with MDC using IHC for desmin stain (brown), B} Minimal increase in thickness of skeletal muscle bundles in PGA only using IHC for desmin stain (brown), C} Abundant nerve fibers in between muscle bundles in PGA seeded with MDC using IHC for S100 stain (brown), D} Minimal increase in nerve fibers in between muscle bundles in PGA only using IHC for S100 stain (brown)

**Figure 3:**
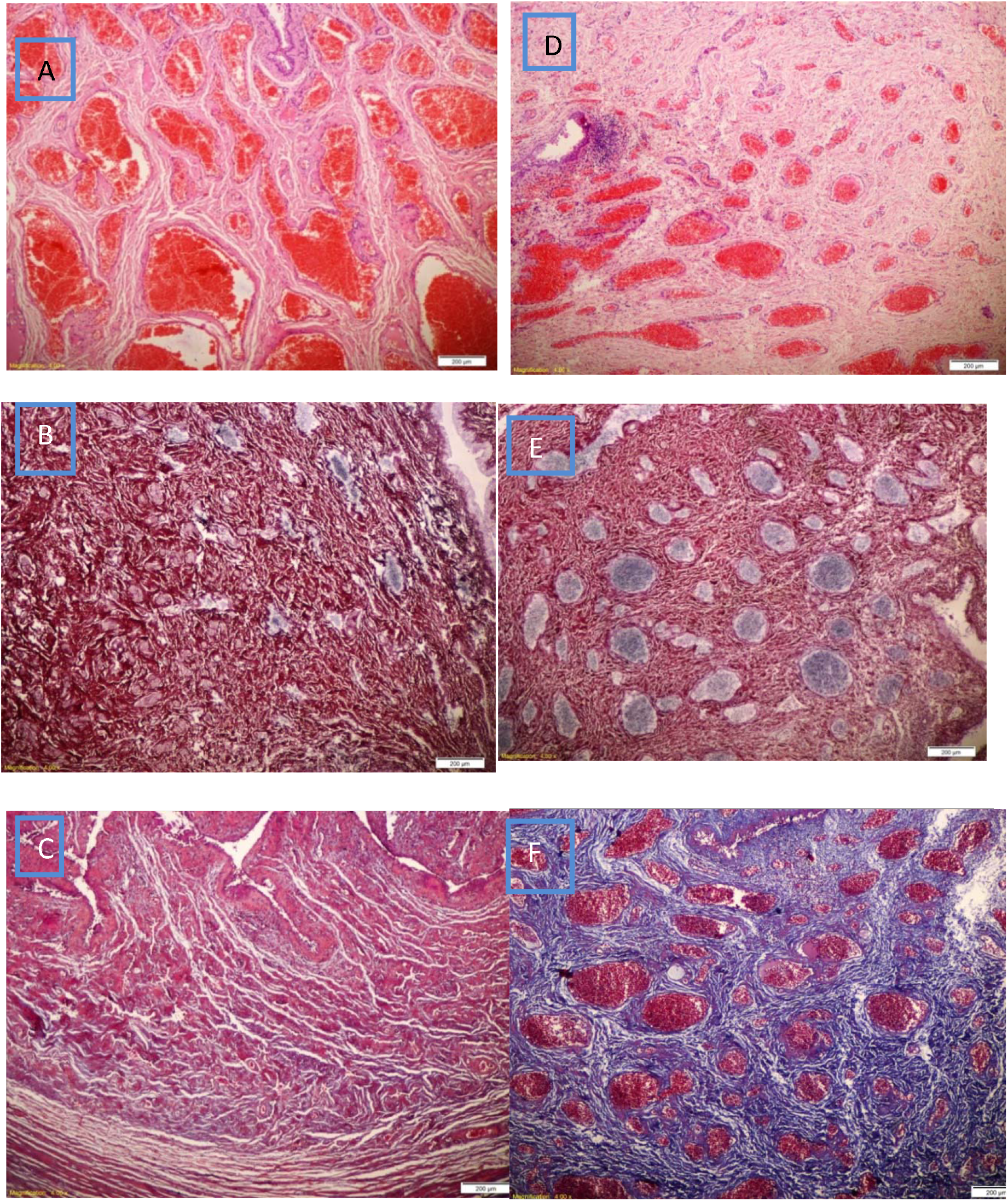
A} Venous plexus is increased in number with marked congestion and mild edema in PGA seeded with MDC, D} Venous plexus is near normal in number with mild congestion with mild inflammatory infiltrate in PGA only, B} Silver stain shows near normal amount of reticular fibers (black) in PGA seeded with MDC, E} Silver stain shows lower amount of reticular fibers (black) in PGA only, C} The lamina propria expressed absent fibrous tissue (blue) in PGA seeded with MDC using Masson trichrome stain, F} The lamina propria showed diffuse moderate fibrous tissue (blue) in PGA only using Masson trichrome stain

## Discussion

Induction of incontinence by urethrolysis with pubourethral ligament injury was our approach of choice. It resulted in disruption of the native support of the urethra and durable loss of urethral resistance^9^. Incontinence was confirmed by lower maximum urethral closure pressure as compared to control dogs, in which no urethrolysis was carried out

Myoblasts have been used as a treatment of stress incontinence in animal model by means of intra-urethral injection^10^. In 2014, a similar approach was used in adult 35 women. Gras et al^11^ described a technique performed largely under local anesthesia/intravenous analgesia. An open muscle biopsy was obtained from vastus lateralis muscle and was “minced” and injected as a suspension in the same session via periurethral route in 35 women; under vaginal US. Cure/improvement was noted in 7% to 63% of the patients.

Other human trials entailed intraurethral injection of muscle-derived stem cells were published, both in adult women^12^ and in children^13^. Although results were promising, some reports turned out to be unfounded^14^.

Another research strategy involved making a structured skeletal muscle tissue in vitro, with the potential of making an all- autologous sling. Many studies evolved to reconstruct skeletal muscle tissue. Some focused on developing a self-organizing tissues, without artificial scaffolds^15^; others preferred seeding cells on a natural or a synthetic biodegradable substrates e.g. collagen matrices ^16^

We preferred Polyglycolic acid as a biodegradable scaffold as it was used by Saxena et al ^17&18^ where myoblasts derived from neonatal rats, Fisher CDF-F344, were seeded onto polyglycolic acid meshes and implanted into the omentum of syngeneic adult Fisher CDF-F344 rats with promising results and comprehensive description of the approach. The advantage of the sling approach is that it applies potentially successful technique^19^ with an added effect of autologous skeletal muscle fibers to the mid-urethra

To our knowledge, no human study involving biodegradable sling seeded with autologous muscle-derived cells was yet reported. Such a study will be of paramount importance, considering that the current treatment options of women with SUI are far from optimum^20^

The question we tried to answer is whether muscle-seeded biodegradable scaffold is any different from plain one? So, we compared absorbable sling to sling seeded with autologous muscle cells.

Cell -seeded sling proved to be more efficacious than plain PGA. Urethral pressure measurement 6 weeks after sling insertion showed that the median increase of urethral closure pressure in the cell-seeded sling was over 40 cm H20 while in the other group where only PGA sling was applied, it was 5.5 cmH20

Histopathological study of harvested urethral segments 4 weeks after insertion of slings proved the persistence of viable skeletal muscle and nerve fibres.

This sling design avoids polypropylene-related adverse events. Application of this technique in humans is easy and promising, based on functional and urodynamic outcome.

One of the shortcomings of our study is the measurement of maximum urethral pressure in female dogs might not be very reproducible. This is probably true according to one study^21^; yet, others^22^ have adopted the same parameter we used

## Conclusion

A biodegradable PGA sling seeded with autologous myoblasts shows evidence of survival 6 weeks after insertion in dogs. The use of skeletal muscle –seeded PGA scaffold is a practical technique with preserved histological differentiation in canine model at short term.

Supplementary figure: Flowchart of the study procedures

## References

1 Nadeau G, Herschorn S.(2014): Management of recurrent stress incontinence following a sling. Curr Urol Rep., 15(8):427

2 Hinoul P, Roovers JP, Ombelet W, Vanspauwen R.(2009): Surgical management of urinary stress incontinence in women: a historical and clinical overview. Eur J Obstet Gynecol Reprod Biol. 145(2):219–225

3 Abrams P, Andersson KE, Birder L et al (2010): Fourth international consultation on incontinence recommendations of the international scientific committee: evaluation and treatment of urinary incontinence, pelvic organ prolapse, and fecal incontinence. Neurourol.Urodyn. 29(1):213–240

4 Daneshgari F, Kong W, Swartz M. (2008): Complications of mid urethral slings: Important outcomes for future clinical trials. J Urol. 180(5):1890–1897

5 Wadie BS, Edwan A, Nabeeh AM. (2005): Autologous fascial sling vs. polypropylene tape at short-term followup: a prospective randomized study. J Urol. 174(3):990–993

6 Blaivas JG, Simma-Chiang V, Gul Z, Dayan L et al (2019): Surgery for stress Urinary Incontinence: Autologous Fascial Sling. Urol Clin North Am. 46(1):41–52

7 Rodríguez LV, Chen S, Jack GS, et al (2005): New objective measures to quantify stress urinary incontinence in a novel durable animal model of intrinsic sphincter deficiency. Am J Physiol Regul Integr Comp Physiol. 288(5):R1332–8

8 Eberli, D, Aboushwareb, T., Soker, S et al (2012): Muscle Precursor Cells for the Restoration of Irreversibly Damaged Sphincter Function, Cell Transplantation. 21 (9), 2089–2098

9 Sajadi KP, Gill BC, Damaser MS. (2010): Neurogenic aspects of stress urinary incontinence. Curr Opin Obstet Gynecol. 22(5):425–429

10 Chancellor MB, Yokoyama T, Tirney S et al (2000): Preliminary results of myoblast injection into the urethra and bladder wall: a possible method for the treatment of stress urinary incontinence and impaired detrusor contractility. Neurourol Urodyn. 19(3):279–287

11 Gräs S, Klarskov N, Lose G. (2014): Intraurethral injection of autologous minced skeletal muscle: a simple surgical treatment for stress urinary incontinence. J Urol. 192(3):850–855

12 Carr LK, Steele D, Steele S, et al (2008): 1-year follow-up of autologous muscle derived stem cell injection pilot study to treat stress urinary incontinence. Int Urogynecol J Pelvic Floor Dysfunct. 19:881–883

13 Kajbafzadeh AM, Elmi A, Payabvash S, et al (2008): Transurethral myoblast injection for treatment of urinary incontinence in children with classic bladder exstrophy. J Urol. 180:1098–1105

14 Kleinert S, Horton R. (2008): Retraction—Autologous myoblasts and fibroblasts versus collagen [corrected] for treatment of stress urinary incontinence in women: A [corrected] randomised controlled trial. Lancet. 372: 789790

15 Dennis RG, KosnikP, Gilbert ME et al (2001): Excitability and contractility of skeletal muscle engineered from primary cultures and cell lines. Am J Physiol Cell Physiol. 280:C288–95

16 KosnikJr P, Dennis RG, Faulkner JA. (2001): Functional development of engineered skeletal muscle from adult and neonatal rats. Tissue Eng. 7(5):573–584

17 Saxena AK, Marler J, Benvenuto M, et al (1999): Skeletal muscle tissue engineering using isolated myoblasts on synthetic biodegradable polymers: preliminary studies. Tissue Eng. 5(6):525–532

18 Saxena AK, Willital GH, Vacanti JP. (2001): Vascularized three dimensional skeletal muscle tissue-engineering. Biomed Mater Eng. 11(4):275–281

19 Richter, H. E., Albo, M. E., Zyczynski, H. M., et al (2010): Retropubic versus transobturator midurethral slings for stress incontinence. N Eng J Med. 362(22): 2066–2076

20 Albo ME, Richter HE, Brubaker L, et al (2007): Burch Colposuspension versus Fascial Sling to Reduce Urinary Stress Incontinence N Engl J Med. 356(21):2143–2155

21 Holt PE (1989): ‘Simultaneous’ urethral pressure profilometry in the bitch: methodology and reproducibility of the technique. Res Vet Sci. 47(1): 110–116

22 Claeys S, de Leval J, Hamaide A. (2010): Transobturator vaginal tape inside out for treatment of urethral sphincter mechanism incompetence: preliminary results in 7 female dogs. Vet Surg. 39(8): 969–979.

